# PDE10A Inhibition Disrupts Lipid Droplet Formation and Sensitizes Macrophages to Ferroptosis after LPS treatment

**DOI:** 10.1101/2025.06.02.657436

**Authors:** Bradford C. Berk, Jinmin Zhang, Chia George Hsu

**Author notes:** Correspondence to: Chia George Hsu, Ph.D.; Co-Correspondence to: Bradford C. Berk, M.D. Ph.D.

## Abstract

Ferroptosis, an iron-dependent form of cell death, plays a key role in various diseases, but its impact on immune cells, particularly macrophages, remains unclear. This study explores how macrophage activation influences susceptibility to ferroptosis, focusing on lipopolysaccharide (LPS) and other inflammatory signals. We found that LPS priming enhanced resistance to ferroptosis in bone marrow-derived macrophages (BMDMs), as shown by reduced morphological changes, lower LDH release, and diminished cell death in real-time assays. Similar effects were observed with Zymosan A and TNF-α. Importantly, LPS-induced ferroptosis resistance was independent of stress response pathways like Nrf2 signaling. Instead, lipid droplet accumulation, driven by LPS, was central to this resistance. PDE10A inhibition reversed LPS-induced ferroptosis and reduced lipid droplet formation. LPS did not confer similar resistance in non-macrophage cell types, underscoring the macrophage-specific nature of this response. These findings highlight potential therapeutic targets for inflammatory diseases.

## Introduction

Ferroptosis is a regulated form of cell death characterized by the accumulation of lipid peroxides to toxic levels, leading to cellular damage and dysfunction[1, 2]. Unlike other forms of cell death, such as apoptosis or necrosis, ferroptosis is driven by iron-dependent oxidative stress and lipid peroxidation, making it a unique and distinct process[1, 2]. Recent studies have highlighted the significance of ferroptosis in various physiological and pathological contexts, including cancer, neurodegeneration, and cardiovascular diseases[3–6]. However, its role in immune cells, particularly macrophages, remains an area of active investigation[7].

Macrophages are critical components of the innate immune system and play a pivotal role in orchestrating inflammatory responses[8]. They are highly versatile cells that can adopt diverse phenotypes, ranging from pro-inflammatory to anti-inflammatory macrophages, depending on the nature of the stimuli they encounter[9]. The ability of macrophages to sense and respond to a wide array of environmental cues, including pathogen-associated molecular patterns (PAMPs) such as lipopolysaccharide (LPS), makes them essential in both defending against infections and in regulating inflammation[10]. However, the response of macrophages to cellular stress, particularly in the context of ferroptosis, is not well understood[7].

Ferroptosis in macrophages is of particular interest because of the potential implications it holds for inflammation, immune regulation, and tissue homeostasis. While macrophages are known to generate reactive oxygen species (ROS) as part of their immune defense mechanisms, excessive oxidative stress can lead to tissue damage[9]. The balance between cell survival and death is crucial in determining the outcome of immune responses, and dysregulation of this balance can contribute to chronic inflammation and autoimmune diseases[11]. Moreover, macrophage ferroptosis may play a role in the resolution of inflammation, tissue repair, and the modulation of adaptive immune responses[9].

Despite its importance, the precise mechanisms underlying ferroptosis in macrophages remain unclear. In cancer cells, transcriptional networks[12], Nrf2 activation[13], and metabolic shifts, such as the Warburg effect[14], have been identified as key regulators of ferroptosis sensitivity. Nrf2, in particular, is activated by oxidative stress to upregulate antioxidant genes and protect cells from ferroptosis[12, 13, 15]. The Warburg effect, which shifts metabolism towards glycolysis, can also enhance resistance to ferroptosis by altering redox balance[14, 16, 17]. Lipid droplets, which store neutral lipids, may serve as a reservoir against ferroptosis via promoting the Warburg effect by inducing FBP1 and FOXO1 [18–20].

In this study, we explored the relationship between inflammation and ferroptosis in macrophages. Phosphodiesterases are enzymes that degrade cAMP and cGMP, key regulators of inflammatory mechanisms, including cytokine and chemokine release, as well as lipid metabolism and lipid droplet formation[21–24]. However, it remains uncertain whether these mechanisms similarly protect macrophages from ferroptotic cell death. Understanding how inflammatory stimuli influence macrophage ferroptosis could offer new therapeutic strategies for diseases marked by excessive cell death or chronic inflammation. We focused on how inflammatory signals modulate macrophage susceptibility to ferroptosis and the molecular pathways involved. This research provides insights into how macrophages balance immune function with cellular survival in inflammatory contexts.

## Results

Lipopolysaccharide (LPS) is widely used to investigate inflammatory macrophages, as it robustly activates these immune cells and triggers inflammatory responses[25, 26]. Exposure to LPS provides valuable insights into the complex molecular processes that underpin inflammation. Glutathione peroxidase 4 (GPX4) plays a crucial role in protecting cells from ferroptosis, a form of regulated cell death, by preventing lipid peroxidation and maintaining cellular redox balance[27, 28].

In this study, we used the nitroisoxazole-containing compound ML210, a prodrug that is converted within cells into a nitrile-oxide electrophile. This electrophile covalently and selectively inhibits GPX4[29]. Treatment with 3 µM ML210 induced ferroptosis in bone marrow-derived macrophages (BMDMs), leading to cell death through this pathway (Fig.1A). To explore the relationship between macrophage activation and ferroptosis, we primed BMDMs with LPS for 24 hours before treating them with ML210. LPS-induced pro-inflammatory macrophages exhibited high resistance to ML210-induced ferroptosis in a dose-dependent manner (Fig.1A-B). This was evident from both morphological changes and the release of lactate dehydrogenase (LDH) (Fig.1A-B). Morphologically, cells undergoing ferroptosis typically exhibit significant swelling, which was observed in non LPS-primed BMDMs treated with ML210. In contrast, LPS-primed macrophages showed less pronounced swelling, indicating their higher resistance to ferroptosis. Additionally, the release of LDH into the culture medium, a well-established marker of cell membrane integrity and cell death, was lower in LPS-primed macrophages, further suggesting their resistance to ferroptosis (Fig.1A-B). To further investigate cell death dynamics, real-time assays using SYTOX™ Green were employed. SYTOX™ Green, a fluorescent dye that selectively stains nucleic acids in cell death, consistently demonstrated that LPS-primed macrophages displayed a significantly lower fluorescence intensity compared to non-primed macrophages (Fig.1C). The combination of morphological observations, LDH release assays, and kinetic cell death assays reinforced these findings, highlighting the protective effects of LPS priming in macrophages against ferroptosis.

**Fig. 1.**
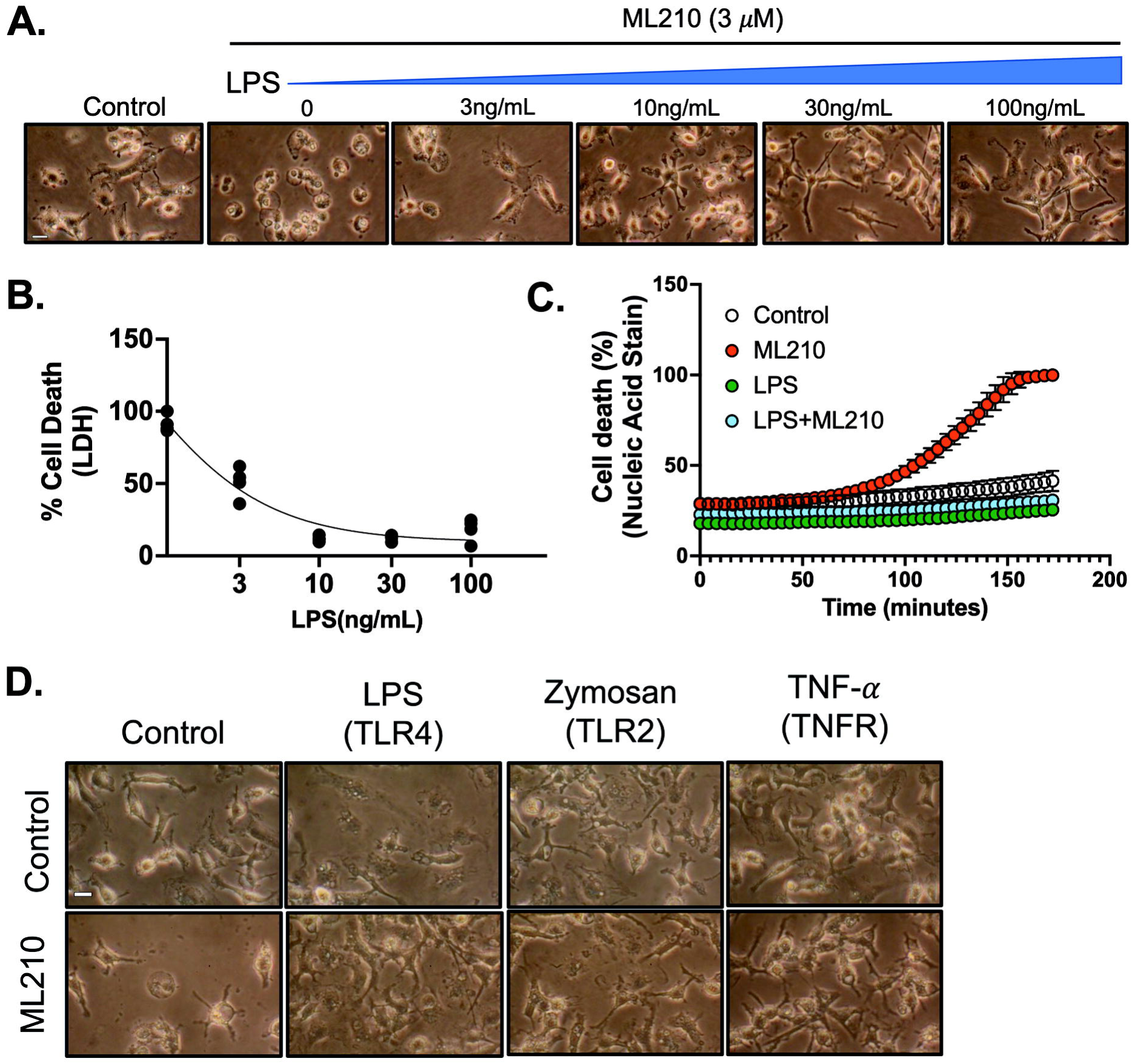
Inflammatory macrophages are highly resistant to ferroptosis. **A-B.** Bone marrow-derived macrophages (BMDMs) were stimulated with or without LPS (3-100 ng/mL) for 24 hours, followed by treatment with ML210 (3 *µ*M) for 12 hours. **A.** Representative images of BMDMs, scale bar= 5 µm. **B.** LDH released was measured in the culture medium, and LDH cytotoxicity was normalized to maximum cell death. **C.** BMDMs were stimulated with LPS (100 ng/mL) for 24 hours, followed by treatment with ML210 (3 *µ*M) for 12 hours. Real-time percentage cell death measurement was performed in the last 3 hours using SYTOXTM green. **D.** BMDMs were stimulated with or without LPS (100 ng/mL), Zymosan (10 *µ*g/mL), TNF-α (20ng/mL) for 24 hours, followed by treatment with ML210 (3 *µ*M) for 12 hours. Representative images of BMDMs, scale bar= 5 µm. Statistical analyses in B and C were conducted using a one-way ANOVA and Bonferroni’s post hoc test. ****P<0.001 between ML210 and ML210+LPS groups. Data are presented as mean±SD. (N=3 experiments).

LPS induces macrophage activation via Toll-like receptor 4 (TLR4), triggering a robust inflammatory response that confers resistance to ferroptosis. However, it remains unclear whether this resistance is specifically driven by LPS-induced signaling or if other inflammatory pathways activated by different Toll-like receptors (TLRs) also contribute to this protective effect. To address this, we utilized Zymosan A, which activates macrophages through TLR2 and TLR6, distinct from the TLR4 pathway activated by LPS. The β-glucan component of Zymosan binds to TLR2, initiating downstream signaling that mirrors LPS activation, including the production of pro-inflammatory cytokines. Our results showed that treatment with Zymosan A similarly protected macrophages from ferroptosis (Fig. 1D), suggesting that TLR2-mediated macrophage activation can confer resistance to ferroptosis independently of LPS signaling. We further examined the role of TNF-α, a key cytokine in the inflammatory response, to determine whether it could induce ferroptosis resistance on its own. TNF-α treatment in macrophages resulted in comparable ferroptosis resistance (Fig. 1D), reinforcing the idea that TNF-α-induced inflammatory pathways can provide protection against ferroptotic cell death. In summary, macrophages treated with Zymosan A and TNF-α demonstrated resistance to ferroptosis similar to that induced by LPS, indicating that the protective effect against ferroptosis is not solely due to LPS but also involves a broader network of inflammatory pathways, including TLR2 and TNF-α signaling.

To gain deeper insights into the effects of LPS on ferroptosis, two additional small molecules were used to induce ferroptosis. Erastin, which inhibits the cystine-glutamate transporter (system XC−)[30], and RSL3, a well-known GPX4 inhibitor, both triggered ferroptosis in macrophages[31]. However, LPS treatment in BMDMs increased resistance to ferroptosis induced by these compounds (Fig. 2A-B). To investigate whether both human and mouse inflammatory macrophages are resistant to ferroptosis, we primed mouse RAW 264.7 macrophages and human THP-1 differentiated macrophages with LPS, followed by treatment with RSL3. This protective effect was observed in both RAW 264.7 macrophages (Fig. 2C) and human THP-1 differentiated macrophages (Fig. 2D). These results show a conserved protective mechanism against ferroptosis in macrophages upon inflammatory activation.

**Fig. 2.**
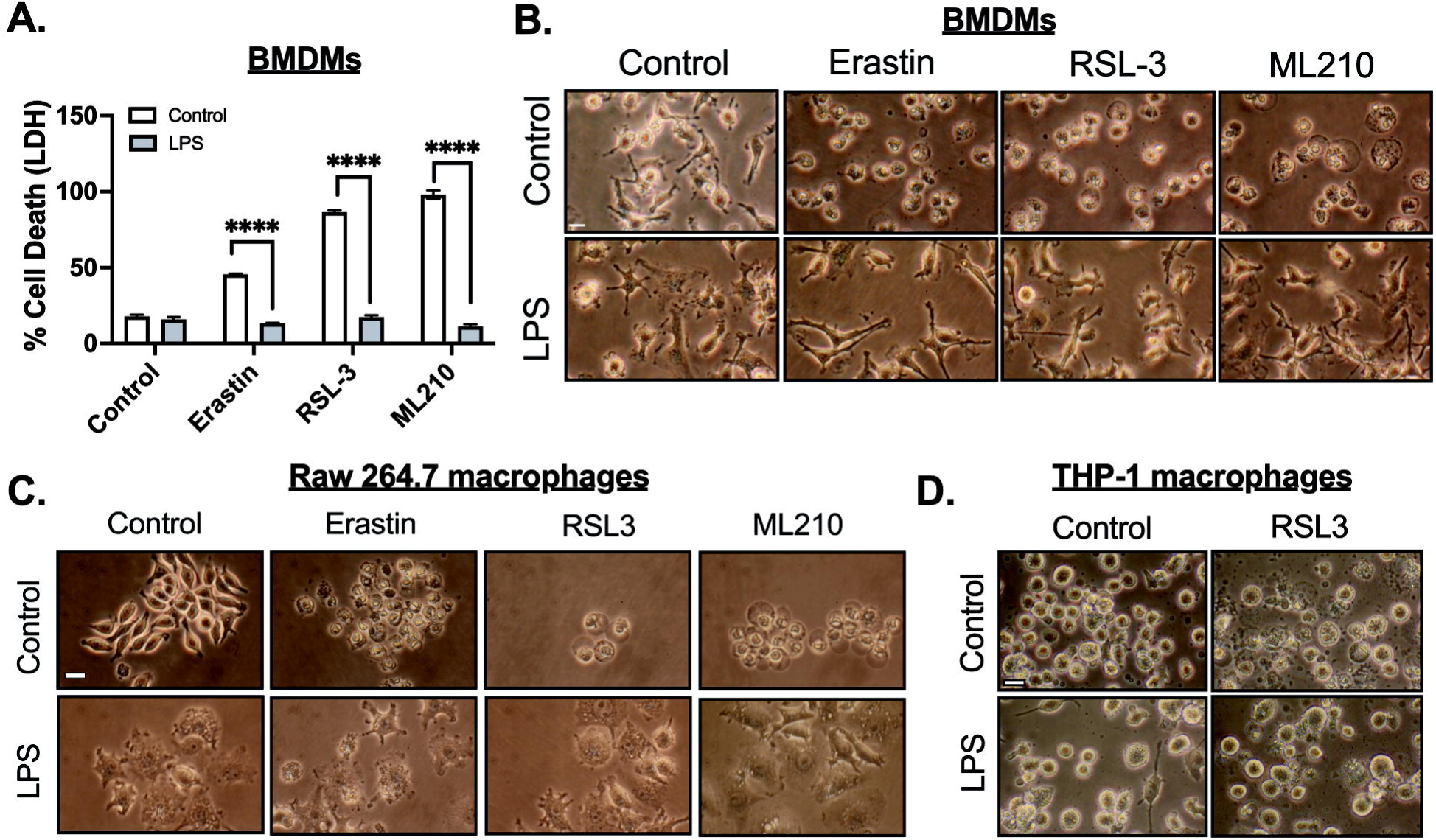
LPS priming enhances ferroptosis resistance in mouse and human macrophages. **A-B.** Bone marrow-derived macrophages (BMDMs) were stimulated with or without LPS (3-100 ng/mL) for 24 hours, followed by treatment with Erastin (10 *µ*M), RSL3 (3*µ*M), or ML210 (3 *µ*M) for 12 hours. **A.** LDH released was measured in the culture medium, and LDH cytotoxicity was normalized to maximum cell death. Bone marrow-derived macrophages (BMDMs) were stimulated with or without LPS (3-100 ng/mL), Zymosan (10 *µ*g/mL), TNF-α (20ng/mL) for 24 hours, followed by treatment with ML210 (3 *µ*M) for 12 hours**. B.** Representative images of BMDMs, scale bar= 5 µm**. C.** Raw 264.7 macrophages were stimulated with or without LPS (100 ng/mL) for 24 hours, followed by treatment with treatment with Erastin (10 *µ*M), RSL3 (3*µ*M), ML210 (3 *µ*M) for 12 hours. Representative images of raw 264.7 cells, scale bar= 5 µm**. D.** Human THP-1 differentiated macrophages were stimulated with or without LPS (100 ng/mL) for 24 hours, followed by treatment with treatment with RSL3 (3*µ*M) for 12 hours. Representative images THP-1 macrophages, scale bar= 5 µm. Statistical analyses in A were conducted using a Two-ways ANOVA and Bonferroni’s post hoc test. ****P<0.001 between control and LPS groups. (N=3 experiments). Bars represent mean ± SD.

Human Umbilical Vein Endothelial Cells (HUVECs) are commonly used to study endothelial function, inflammatory responses, and vascular integrity. LPS treatment is known to induce inflammation in endothelial cells, as evidenced by the increased expression of pro-inflammatory markers such as Monocyte Chemoattractant Protein 1 (MCP-1), Interleukin-1 beta (IL-1β), and Vascular Cell Adhesion Molecule 1 (VCAM-1). (Fig. 3A). These markers are typically associated with endothelial activation during inflammatory responses, such as immune cell recruitment to sites of injury. Despite the robust inflammatory response induced by LPS, the lack of protection against ML210-induced ferroptosis (Fig. 3B) in HUVECs suggests that endothelial cells do not exhibit a ferroptosis-resistant phenotype upon LPS activation. A549 cells, a human alveolar basal epithelial cell line, are commonly used as a model to study lung cancer and epithelial responses to stressors. Similar to HUVECs, LPS treatment in A549 cells induced the expected inflammatory markers, but it did not provide protection against RSL3-induced ferroptosis (Fig. 3C). C2C12 myoblasts, which can differentiate into myotubes and serve as a model for skeletal muscle biology, have been shown in previous studies to be affected by LPS-induced muscle atrophy[32–34]. Despite this well-documented inflammatory response in myotubes, LPS treatment did not protect against RSL3-induced ferroptosis (Fig. 3D). These results highlight the cell-type specificity of LPS-induced resistance to ferroptosis. While LPS protects macrophages from ferroptosis, it does not confer the same protection in endothelial cells, epithelial cells, or myotubes. This specificity could have important implications for understanding immune cell survival in inflammation and for developing therapies targeting ferroptosis in various diseases.

**Fig. 3.**
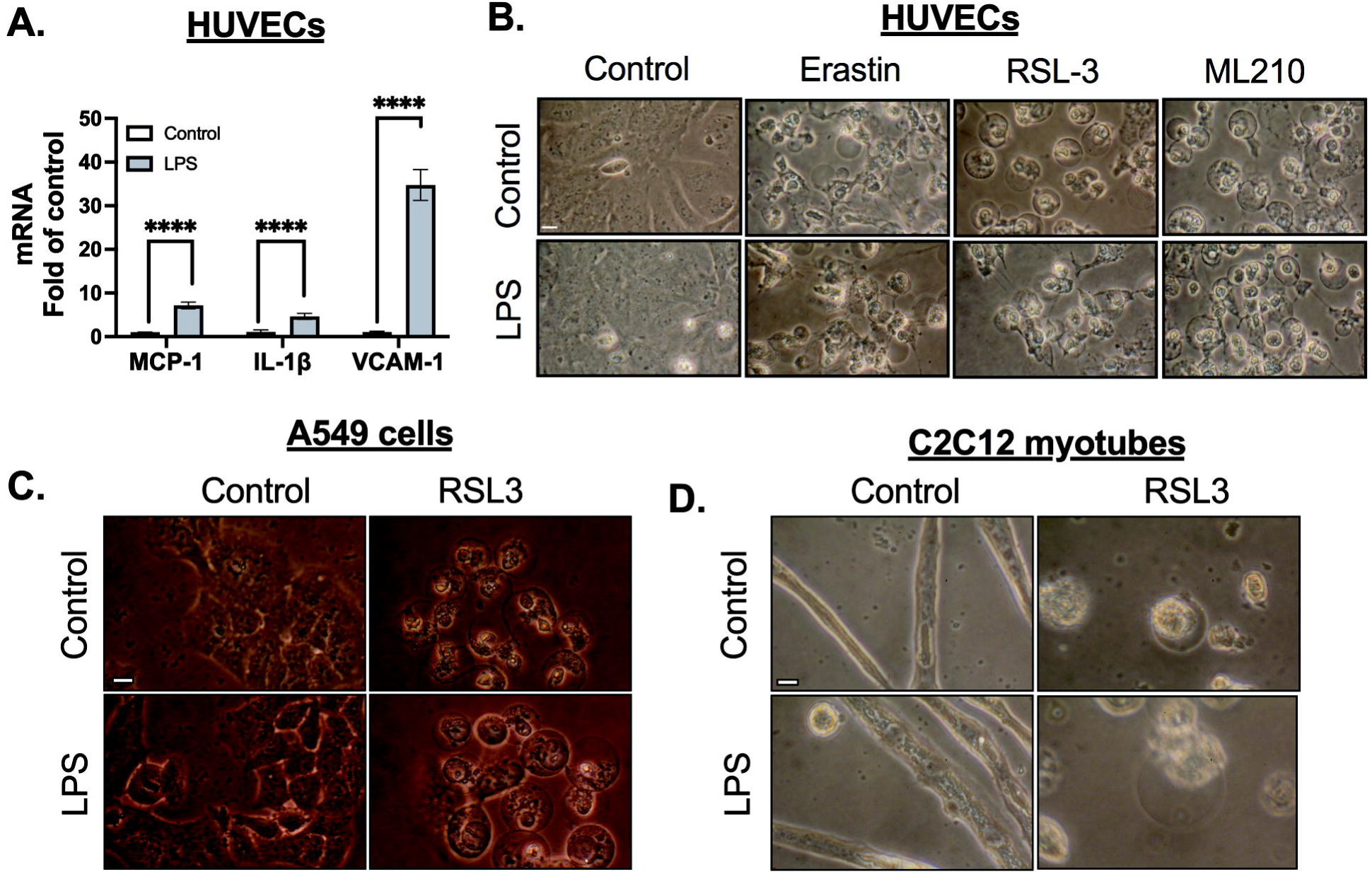
Inflammatory stimulation increases ferroptosis resistance in macrophages. **A.** Human umbilical vein endothelial cells (HUVECs) were stimulated with LPS (100 ng/mL) for 24 hours. Gene expression was analyzed by real-time PCR. MCP-1, IL-1β, and VCAM-1 gene expressions were measured after normalizing to β-actin expression. **B.** In separate experiments, HUVECs were stimulated with LPS (100 ng/mL) for 24 hours followed by treatment with Erastin (10 *µ*M), RSL3 (3*µ*M), or ML210 (3 *µ*M) for 12 hours. Representative images of HUVECs, scale bar= 5 µm. **C.** A549 cells were stimulated with LPS (100 ng/mL) for 24 hours followed by treatment with RSL3 (3*µ*M) for 12 hours. Representative images of A549 cells, scale bar= 5 µm. **D.** C2C12 myotubes were stimulated with LPS (100 ng/mL) for 24 hours followed by treatment with RSL3 (3*µ*M) for 12 hours. Representative images of A549 cells, scale bar= 5 µm. Statistical analyses in A were conducted using a one-way ANOVA and Bonferroni’s post hoc test. ****P<0.001 between control and LPS groups. (N=3 experiments). Bars represent mean ± SD.

To investigate the mechanism of ferroptosis resistance in inflammatory macrophages, we first focused on the role of Nrf2 (Nuclear Factor Erythroid 2-Related Factor 2). Nrf2 is widely recognized for its cytoprotective effects, with multiple lines of evidence supporting its role in protecting cells from ferroptosis[12, 13, 15]. Nrf2 is a transcription factor that plays a crucial role in cellular defense against oxidative stress and in maintaining redox homeostasis[13]. To investigate the potential involvement of Nrf2 in LPS-induced resistance to ferroptosis, we first examined the impact of LPS on Nrf2-regulated genes, such as glutamate-cysteine ligase catalytic subunit (GCLC) and solute carrier family 40 member 1 (SLC40A1). Notably, LPS treatment did not elevate the expression of these genes across a wide range of concentrations or time points (Fig. 4A-D). Similarly, the downstream target of Nrf2, GPX4, which plays a key role in defending against lipid peroxidation, did not show increased gene or protein expression in response to LPS (Fig. 4E-F). Prior research has demonstrated that 4-hydroxynonenal (HNE) promotes antioxidant expression by activating Nrf2 signaling[25, 35]. This activation occurs through the disruption of the Keap1–Nrf2 association, preventing Nrf2 degradation. To assess the role of Nrf2 in protecting macrophages from ferroptosis, we primed macrophages with HNE to activate Nrf2, followed by treatment with ML210. However, HNE showed no significant effect on protecting macrophages from ferroptosis (Fig. 4G). To determine whether Nrf2 activation is essential for LPS-induced resistance to ferroptosis, we used the specific Nrf2 inhibitor ML385 to block Nrf2 signaling. Again, even in the presence of ML385, LPS still prevented cell death (Fig. 4G). These collective findings suggest that the protective effect of LPS against ferroptosis occurs independently of the Nrf2 pathway.

**Fig. 4.**
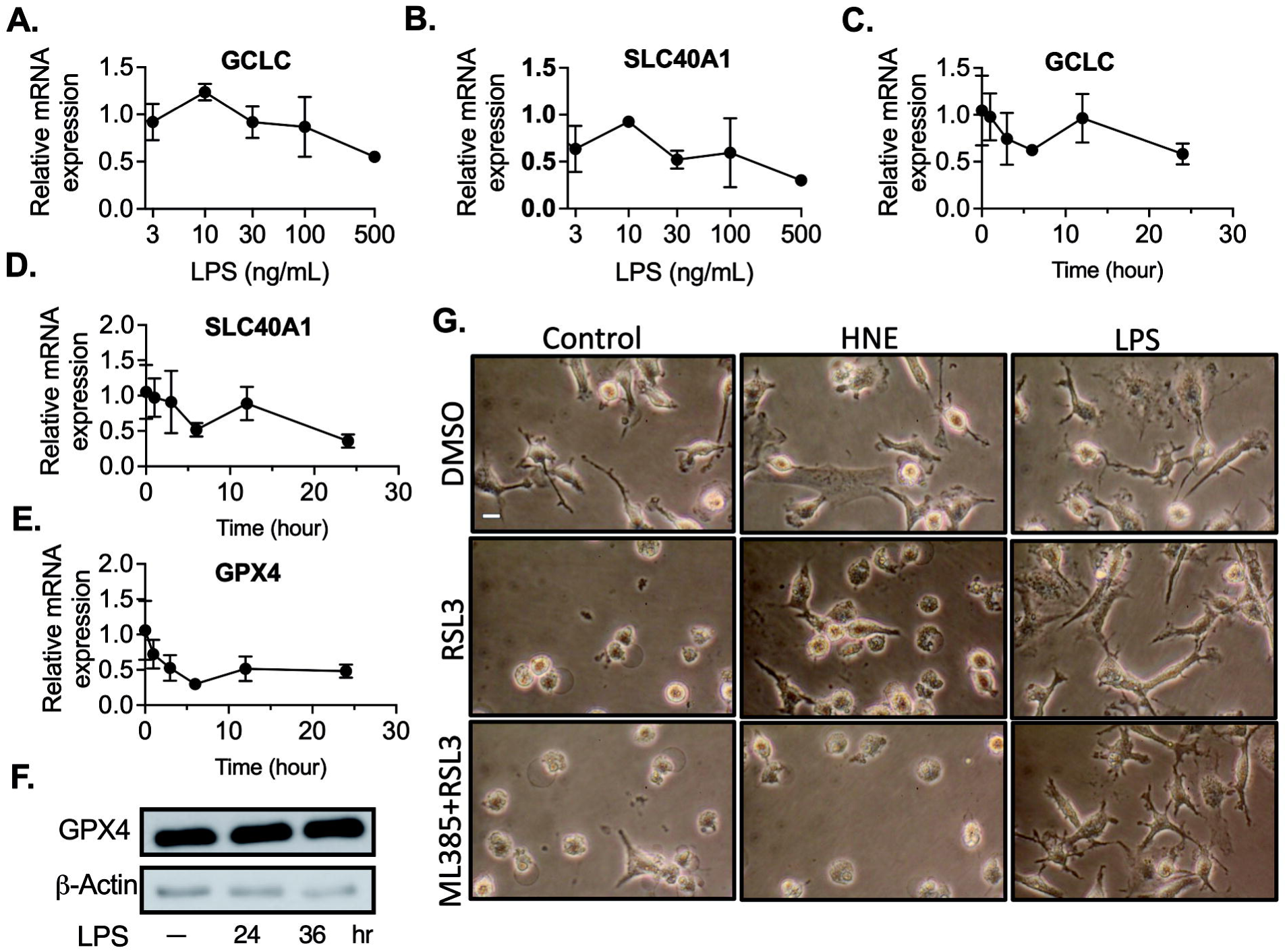
LPS enhances ferroptosis resistance independent of NRF2 signaling in macrophages. **A-B.** Bone marrow-derived macrophages (BMDMs) were stimulated with or without LPS (3-500 ng/mL) for 24 hours. Gene expression was analyzed using Real-Time PCR after normalization to β-actin expression for **A.** GCLC and **B.** SLC40A1. C-E. In the next set, BMDMs were stimulated with LPS for 1-24 hours, and gene expression was analyzed by Real-Time PCR after normalization to β-actin expression for **C.** GCLC**, D.** SLC40A1, and **E.** GPX4. F. BMDMs were stimulated with LPS for either 24 or 36 hours, and protein expression of GPX4 was analyzed by western blot. **G.** BMDMs were stimulated with 4-hydroxynonenal (HNE, 3 *µ*M), LPS with or without ML385 for 24 hours, followed by treatment with RSL3 (3 *µ*M) for 12 hours. Representative images of BMDMs are provided, scale bar= 5 µm. Data are presented as mean±SD. (N=3 experiments).

Several lines of evidence support the idea that a metabolic transition from mitochondrial respiration to glycolysis plays a role in protecting cells from ferroptosis[14, 16, 17]. This metabolic adaptation, commonly known as the glycolytic shift, can impact cellular redox balance and reduce the vulnerability of cells to ferroptotic cell death. In our investigation of the impact of LPS on macrophage metabolic reprogramming, we measured oxygen consumption and extracellular acidification rates under glycolytic stress using a Seahorse analyzer. As expected, LPS priming resulted in a decrease in oxygen consumption, along with an increase in maximal glycolysis and glycolytic capacity, indicating enhanced metabolic activity and a shift toward glycolytic pathways (Fig. 5A-D). Given that glucose is a key energy source for cells and its metabolism is essential for cellular function, we aimed to investigate how glycolysis-dependent metabolism influences LPS-induced ferroptosis. Macrophages were primed with LPS in both high-and low-glucose DMEM, followed by treatment with ML210. Importantly, the removal of extracellular glucose did not diminish the protective effects of LPS (Fig. 5E). These results suggest that LPS-induced metabolic reprogramming and ferroptosis resistance may involve mechanisms beyond glucose availability.

**Fig. 5.**
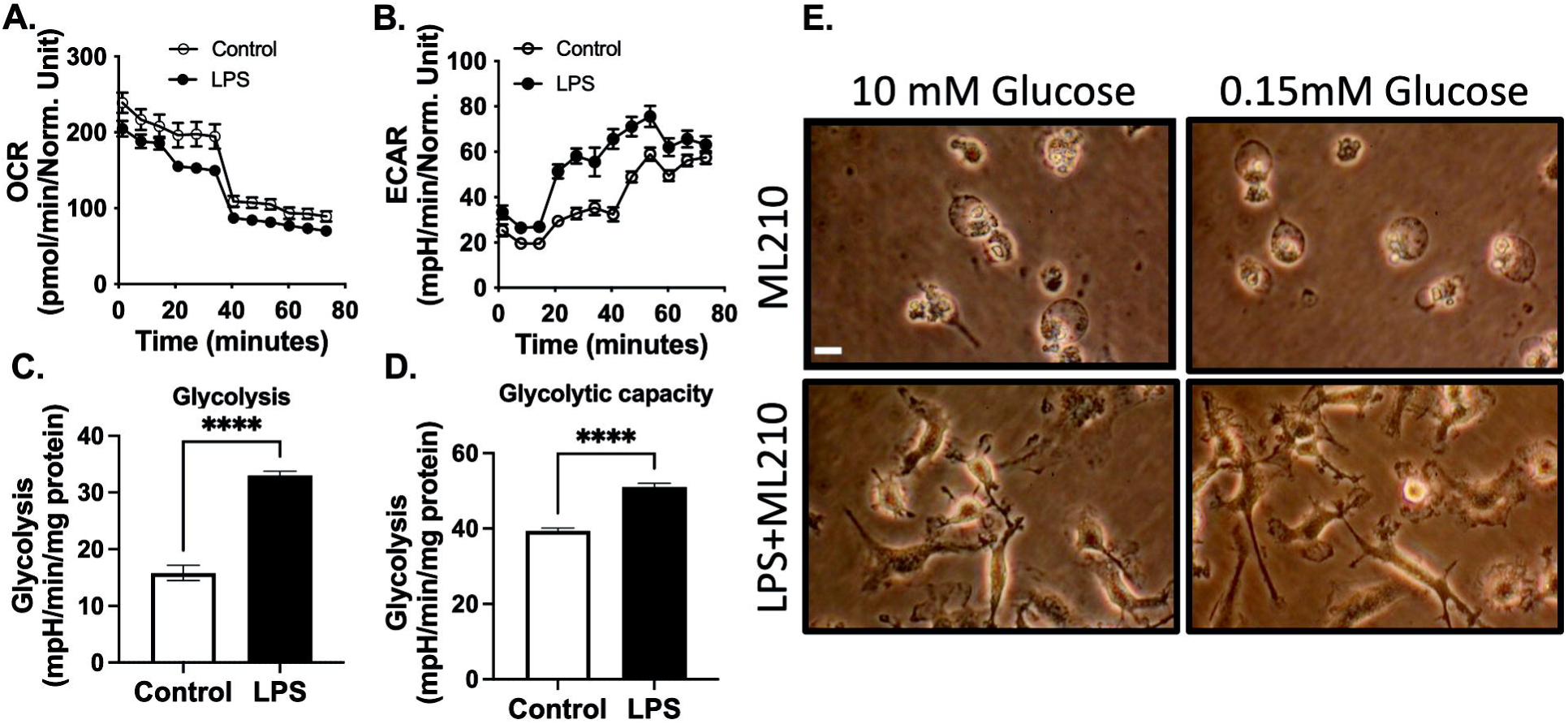
LPS enhances ferroptosis resistance independent of glycolysis. **A-D.** Bone marrow-derived macrophages (BMDMs) underwent a glycolytic stress test using a Seahorse XF-analyzer. Representative Seahorse plots are presented after the injection of glucose (10mM), oligomycin (1 μg/ml), and 2-DG (100mM) in final concentration. The plots illustrate **A.** Oxygen Consumption Rate (OCR) and **B.** Extracellular Acidification Rate (ECAR). **C.** Quantification of glycolysis, defined as the highest rate after glucose stimulation. **D.** Quantification of glycolytic capacity, defined as the highest ECAR after oligomycin treatment. **E.** In separate experiments, BMDMs were stimulated with or without LPS (100 ng/mL) for 24 hours in 10 mM or 0.15 mM glucose DMEM medium, followed by treatment with ML210 (3 *µ*M) for 12 hours. Representative images of BMDMs are provided, scale bar= 5 µm. The data in the figures represent mean±SD (N=3 experiments).

Lipid droplets serve as storage depots for neutral lipids, including fatty acids, and have been proposed to act as a physical barrier, preventing the propagation of lipid peroxidation and subsequent membrane rupture associated with ferroptosis[18–20]. Recent studies have shown that exogenous oleic acid increases ferroptosis resistance in epithelial cells[20, 36]. However, the effects of oleic acid in macrophages have never been explored. THP-1 differentiated macrophages were treated with oleic acid for 24 hours. We observed accumulation of lipid droplet on the cell surface (Fig. 6A). Furthermore, exogenous oleic acid suppressed ML210-induced ferroptosis (Fig. 6B). To examine the potential role of LPS-induced lipid droplet formation, we performed immunofluorescence staining by using BODIPY (493/503). LPS treatment showed strong accumulation of lipid droplet formation (Fig. 6C). Inhibition of phosphodiesterases (PDEs), enzymes that degrade cAMP and cGMP, can lead to the accumulation of these cyclic nucleotides. More specifically, cAMP signaling has been linked to the regulation of lipolysis and lipid metabolism[21–24]. Alterations in lipid metabolism could potentially impact the availability of polyunsaturated fatty acids (PUFAs), which are substrates for lipid peroxidation in ferroptosis. Phosphodiesterase 10A (PDE10A) inhibition has been associated with macrophage activation and inflammation[26, 37]. To determine the role of PDE10A in ferroptosis, we treated BMDMs with specific PDE10A inhibitors, MP10 and TP-10, along with LPS. In both cases, PDE10A inhibition reversed LPS-induced ferroptosis resistance and reduced lipid droplet formation (Fig. 6C, D). These findings suggest that LPS-induced resistance to ferroptosis may be attributed to the formation of lipid droplets. The inhibition of PDE10A, which affects lipid metabolism and macrophage activation, further supports the importance of lipid droplet formation in modulating ferroptosis resistance. These results highlight the complex interplay between lipid metabolism, lipid droplet formation, and ferroptosis resistance, offering potential therapeutic targets for modulating ferroptosis in inflammatory conditions.

**Fig. 6.**
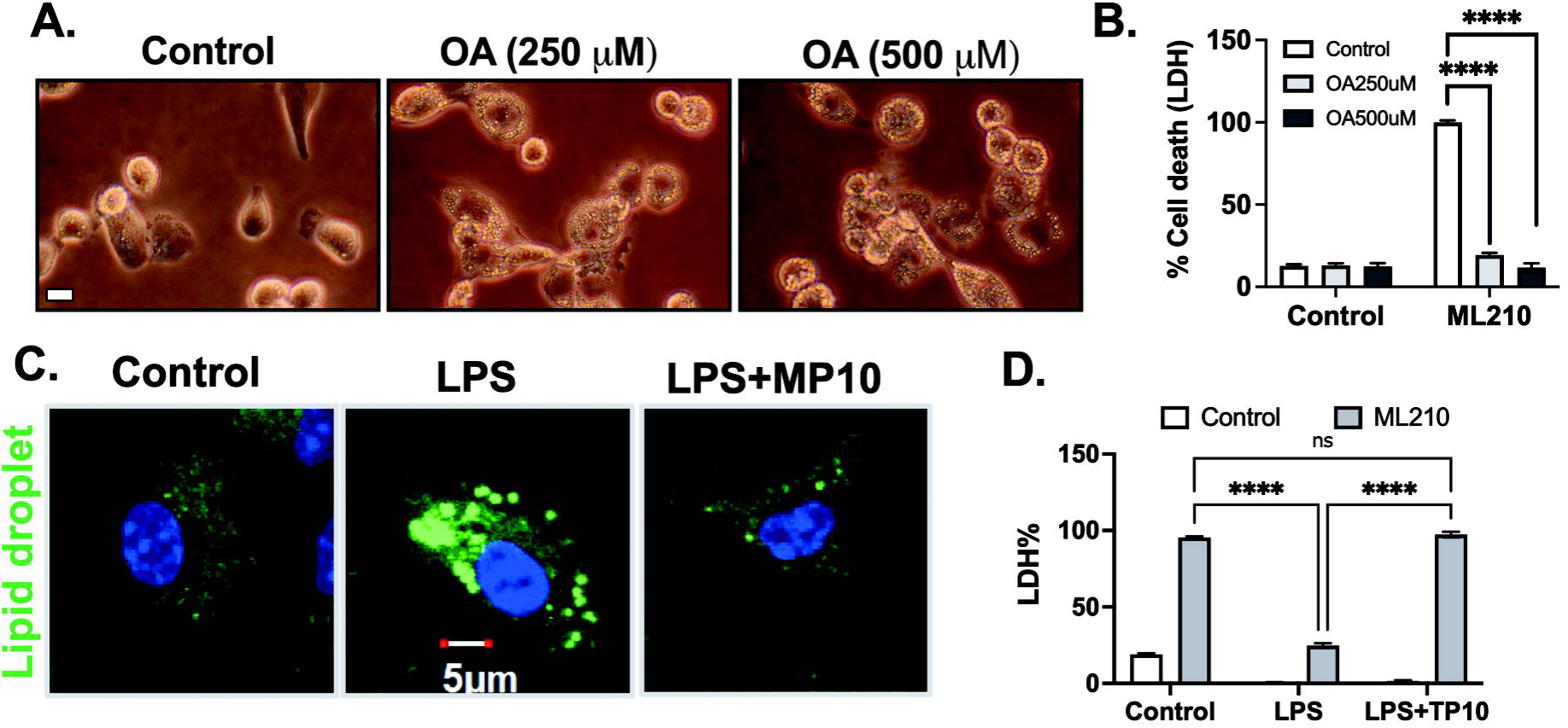
PDE10A inhibition sensitizes ferroptosis and inhibits pyroptosis in activated macrophages. **A.** THP-1-differentiated macrophages were stimulated with or without Oleic acid (250-500 *µ*M) for 24 hours. Representative images of THP-1 macrophages are provided, scale bar= 5 µm. **B.** In separate experiments, THP-1-differentiated macrophages were stimulated with or without Oleic acid (250-500 *µ*M) for 24 hours, followed by treatment with ML210 (3 *µ*M) for 12 hours. LDH release was measured in the culture medium, and LDH cytotoxicity was normalized to maximum cell death. **C.** Bone marrow-derived macrophages (BMDMs) were stimulated with LPS (3-100 ng/mL) or LPS+MP10 (5 *µ*M, PDE10A inhibitor) for 24 hours. Lipid droplets were measured by BODIPY (493/503) immunofluorescence (scale bar: 5 µm), DAPI (blue), and lipid droplet (green). **D.** In separate experiments, BMDMs were stimulated with LPS (100 ng/mL) or LPS+TP10 (5 *µ*M, PDE10A inhibitor) for 24 hours, followed by treatment with ML210 (3 *µ*M) for 12 hours. LDH release was measured in the culture medium, and LDH cytotoxicity was normalized to maximum cell death. Statistical analyses in B, and D were conducted using a two-way ANOVA and Bonferroni’s post hoc test. *P<0.05, ****P<0.001 among groups. (N=3 experiments). Bars represent mean ± SD.

## Discussion

This study investigated the intricate interplay between the inflammatory response and ferroptosis resistance in macrophages. We found that LPS induced a specific and robust resistance to ferroptosis in these cells. Macrophages treated with Zymosan A and TNF-α showed a similar resistance to ferroptosis as that induced by LPS, highlighting that the ferroptosis resistance is not solely due to LPS treatment but is instead a broader response linked to inflammation. The findings provide valuable insights into the complex relationship between the inflammatory response and ferroptosis sensitivity in macrophages. Notably, LPS-mediated protection against ferroptosis is independent of Nrf2 signaling and glycolytic metabolism shifts. Our study also reveals a novel regulatory role for PDE10A in lipid droplet formation and the sensitivity to ferroptosis in macrophages.

In this study, we observed that inflammatory macrophages were highly resistant to ferroptosis across several different macrophage types, including both human and mouse macrophages. This resistance was evident with various ferroptosis-inducing stimuli, such as ML210, erastin, and RSL3. The diverse stimuli—LPS, zymosan, and TNF-α—activate macrophages through distinct receptors but ultimately converge to induce ferroptosis resistance, suggesting the existence of common downstream pathways leading to this shared outcome. This underscores the complex nature of macrophage activation and its influence on cellular responses, particularly in the context of ferroptotic cell death. Given that TNF-α signaling appears to confer resistance to ferroptosis, it may play a crucial role in immune responses during inflammatory diseases. Further research is needed to explore whether chronic inflammatory signaling has lasting effects on ferroptosis resistance in macrophages, or if this resistance itself contributes to the persistence of chronic inflammation in human diseases. Moreover, investigating the intricate signaling pathways triggered by these stimuli in vitro, in vivo, and in human tissues will provide essential insights into the regulatory mechanisms underlying ferroptosis resistance in human disease.

Despite macrophages, endothelial cells, epithelial cells, and muscle cells recognizing LPS through TLR4, the downstream signaling pathways triggered by LPS binding to TLR4 appear to differ between macrophages and other cell types. This difference in signaling pathways likely contributes to the unique cellular responses observed in macrophage ferroptosis. Exploring the molecular and signaling differences between macrophages and other cells could provide valuable insights into the macrophage-specific resistance to ferroptosis in future studies. Additionally, while this study focuses on macrophage ferroptosis, other immune cells, such as neutrophils, B cells, and T cells, may also exhibit resistance to ferroptosis following activation[38, 39]. Therefore, future studies should expand the analysis to include a broader range of immune cell populations following LPS challenge.

Macrophages generate substantial amounts of reactive oxygen species (ROS) with antimicrobial properties against various pathogens. However, the mechanisms by which macrophages sense intracellular ROS levels and protect themselves from oxidative stress-induced damage remain incompletely understood. The involvement of NRF2 signaling as an oxidative stress response mechanism in macrophages suggests its potential role in protecting cells from oxidative stress in challenging environments during infections. Our previous research demonstrated that HNE treatment induces Nrf2-dependent expression of GCLC and SLC40A1, key genes that may help prevent oxidative stress[25]. However, HNE priming does not protect macrophages from ferroptosis. Additionally, the Nrf2 inhibitor ML385 does not sensitize macrophages to ferroptosis following LPS treatment. While Nrf2 activation has been shown to have a protective role in cancer cells[40–43], cardiomyocytes[44], and epithelial cells[45], further studies are needed to clarify the specific role of Nrf2 in macrophages in relation to ferroptosis.

Previous studies have shown that exogenous monounsaturated fatty acids (MUFAs) effectively inhibit ferroptosis in cancer cells by suppressing the accumulation of toxic lipid reactive oxygen species (ROS), particularly at the plasma membrane and endoplasmic reticulum[20, 36, 46]. This protective mechanism involves reducing phospholipids containing oxidizable polyunsaturated fatty acids. In the presence of MUFAs, lipid droplet formation plays a key role by preventing the buildup of polyunsaturated fatty acids in cellular membranes, thereby averting lipid peroxidation associated with ferroptosis[20, 36, 46]. The regulation of lipid droplets and lipid composition is crucial in determining the fate of cancer cells, particularly in conferring resistance to ferroptosis[47, 48]. In our study, LPS treatment resulted in a marked increase in lipid droplet formation, suggesting a potential link between LPS-induced ferroptosis resistance and lipid droplet accumulation. Moreover, our finding that PDE10A inhibition reversed LPS-induced ferroptosis and reduced lipid droplet formation offers a promising therapeutic target. PDE10A inhibitors, which have been explored for other clinical uses, may provide a novel approach to modulate ferroptosis in macrophages[49, 50]. Given the role of lipid droplets in ferroptosis resistance, modulating lipid metabolism through PDE10A inhibition may enhance cell survival during inflammatory responses. Further investigation into PDE10A’s molecular mechanisms in lipid metabolism and ferroptosis may reveal new therapeutic opportunities for immune modulation, particularly for diseases involving dysregulated cell death, such as sepsis or chronic inflammation.

Our study focused on macrophage resistance to ferroptosis, but the factors that sensitize macrophages to ferroptosis are still not well understood. Recent research has identified polyunsaturated phosphatidylcholine species (PC-PUFA2s) as key initiators of ferroptosis in cancer cells[20, 36, 46]. Macrophages in different microenvironments may express varying levels of PC-PUFA2s or PE-PUFA, influencing their susceptibility to ferroptosis. Whether pro-inflammatory or anti-inflammatory macrophages exhibit significant differences in PC-PUFA2 levels remains to be investigated.

In summary, this study explored the resistance of inflammatory macrophages to ferroptosis, triggered by LPS, zymosan A, and TNF-α. We demonstrate that this resistance is not reliant on Nrf2 signaling or glycolytic shifts but is linked to lipid droplet accumulation. Additionally, inhibition of PDE10A reverses ferroptosis resistance and lipid droplet formation. These findings provide insights into macrophage-specific ferroptosis resistance, highlighting potential therapeutic targets for inflammatory diseases.

## Materials and Methods

### Bone marrow progenitor cell isolation and bone marrow-derived macrophage (BMDM) differentiation

BMDMs preparation was performed as previously described [51]. L929 conditioned media which contains the macrophage growth factor M-CSF, was prepared by culturing L929 cells (ATCC) in complete DMEM (Thermo Fisher Scientific, MT10013CV) supplemented with 10% FBS, and 1% penicillin and streptomycin for 10 days at 37°C, 5% CO_2_. The L929 conditioned media, was collected, filtered (Vacuum Filter/Storage Bottle System, Corning, 431153), and stored at −80 °C until required. For isolation of BMDMs, tibias and femurs were removed from both male and female mice and flushed with media using a 26-gauge needle. Bone marrow was collected at 500 x g for 2 min at 4 °C, resuspended with complete DMEM medium and filtered through a 70-μm cell strainer (VWR international, 10199-657). Bone marrow progenitor cells were cultured in 100 mm dishes for 6-7 days in 70% complete DMEM medium and 30% L929-conditioned medium. Fresh medium (5 mL) was added on day 3. BMDMs were collected by scraping in cold PBS containing EDTA (1 mM). After centrifugation, BMDMs were seeded into 12-well plates at a density of 1.6 × 10^5^ cells/well in DMEM and incubated overnight before use.

### Raw 264.7 macrophages

Raw 264.7 cells were cultured in 12-well plates at a density of 1.6 × 10 cells/well in DMEM containing 10% FBS and 1% penicillin-streptomycin, at 37°C in a 5% CO₂ incubator. Before the experiment, the cells were washed twice and incubated in serum- and antibiotic-free DMEM.

### THP-1 macrophage differentiation

THP-1 differentiated macrophages were prepared as previously described[52]. Human THP-1 monocytes were differentiated into macrophages by 24 hr incubation with 100 nM PMA (Sigma-Aldrich) in complete RPMI medium at 1.6 × 10^5^ cells/well in 12-well plates. Cells were washed twice with 1x PBS and incubated with complete RPMI medium without PMA for 24 hr before experiment.

### HUVEC isolation

HUVEC (Human Umbilical Vein Endothelial Cells) isolation from human umbilical cord was performed as previously described [53]. Four donated umbilical cord veins were digested with collagenase type I (1 mg/mL) for 20 minutes at 37°C, followed by centrifugation for approximately 10 minutes at 1000 rpm and 4°C. The cells were then cultured in complete medium (M200 with 5% FBS, 1% streptomycin/penicillin, and 2% LSGS (Gibco™, S00310)). Cells were frozen at passages 2-3. Prior to the experiment, the cells were allowed to reach confluence, washed twice, and incubated in serum- and antibiotic-free DMEM.

### A549 cells

A549 cells were cultured in 12-well plates at a density of 1.6 × 10 cells/well in DMEM containing 10% FBS and 1% penicillin-streptomycin, at 37°C in a 5% CO₂ incubator. Before the experiment, the cells were washed twice and incubated in serum- and antibiotic-free DMEM.

### C2C12 myotubes

C2C12 myoblasts were cultured in growth medium consisting of DMEM with 10% FBS, 1% penicillin and streptomycin, and maintained in a humidified incubator at 37°C with 5% CO2. The cells were seeded at a density of 4.4 × 10³ cells/cm² in 12-well plates and allowed to proliferate for 2 days, reaching 60-80% confluence. At this point, they were switched to differentiation medium (DMEM supplemented with 2% horse serum (Thermo Fisher Scientific, 16050122) and antibiotics), with media changes occurring every 48 hours. Myoblasts typically fused into myotubes within 4 days[54, 55]. For treatment, the cultures were washed twice with serum- and antibiotic-free DMEM.

### Inflammatory stimulation

Macrophage cultures were rinsed twice with serum- and antibiotic-free medium. Cells were then exposed to LPS (3–100 ng/mL), Zymosan (10 µg/mL), or TNF-α (20 ng/mL) for 24 hours in media. Endothelial cells C2C12 myotubes, A549 cells, and human umbilical vein endothelial cells (HUVECs) were stimulated with LPS (100 ng/mL) for 24 hours.

### Ferroptosis activation

BMDMs, THP-1 differentiated macrophages or HUVECs were treated with Erastin (10 *µ*M), RSL3 (3*µ*M), or ML210 (3 *µ*M) for 12-24 hours in serum and antibiotic-free medium.

### Fatty acid treatment

THP-1-differentiated macrophages were stimulated with or without Oleic acid (250-500 *µ*M) (Cayman, 90260) for 24 hours, followed by treatment with ML210 (3 *µ*M) for 12 hours.

### PDE10A inhibition

Macrophage cultures were rinsed twice with serum- and antibiotic-free medium. Cells were then exposed to LPS (100 ng/mL) with or without two different PDE10A inhibitors (MP-10 at 5 µM, TP-10 at 5 µM) for 24 hours in media.

### Cell morphology

Micrographs of cell cultures were obtained under phase contrast illumination using 40X objective (Leica) prior to cell lysis. Multiple random fields were captured for each well.

### Cell death LDH assay

Culture supernatants were collected and centrifuged at 500 × g for 5 min to remove cellular debris. LDH measurement was performed with the CyQUANT™ LDH cytotoxicity assay kit (Thermo Fisher Scientific, C20301) according to the manufacturer’s instructions. Data were plotted normalizing the O.D. value obtained in wells treated with Triton X-100 (0.1%) as 100%.

### Cell death by SYTOX^TM^ green

BMDMs were seeded in 96-well plates (2×10^4^ cells/well) one day before the experiments. Cells were washed twice and incubated with LPS (100 ng/mL) in XF based medium (Agilent, 103334-100) supplemented with 4.5 g/L glucose, 2 mM glutamine, 1 mM sodium pyruvate, and 1 mM HEPES buffer at final pH7.4 for 3 hr. After 24 hr LPS stimulation, SYTOX Green (final concentration 1µM) (Thermo Fisher Scientific, S7020) was added together with ML-210 (3 µM). Cells were incubated at 36°C for 8 hours in a non-CO₂ incubator. Afterward, fluorescence signals (excitation wavelength: 485 nm, emission wavelength: 550 nm) were analyzed using a FLUOstar OPTIMA plate reader (BMG Labtech) at 36°C for 120 minutes. The percentage cell death was calculated by normalizing fluorescence signals from cells treated with Triton X-100 (0.1%).

### Lipid droplet assay

Cells were fixed in 4% paraformaldehyde for 10 minutes, washed three times with 1x PBS, and mounted with Fluoromount-G-DAPI (Thermo Fisher Scientific, 00-4959-52).

Immunofluorescence of lipid droplets and nuclei was analyzed by confocal microscopy, using BODIPY (493/503) (Thermo Fisher Scientific, D3922) for lipid droplets (green) and DAPI (blue) for nuclei.

### Metabolism analysis

Glycolytic and mitochondrial activity of macrophages was assessed using a Seahorse XF 96 Analyzer (Seahorse Bioscience). BMDMs were seeded at a density of 1 × 10 cells/well in 100 µL DMEM culture medium and incubated overnight. The following day, cells were treated with or without LPS for 24 hours. After treatment, cells were washed and incubated in Seahorse XF base medium (Agilent, 103334-100) containing no glucose, 1X Glutamax (Thermo Fisher Scientific, 1798324), and 1X sodium pyruvate (Thermo Fisher Scientific, 11360070) for 1 hr in a CO₂-free incubator. Oxygen consumption rate (OCR) and extracellular acidification rate (ECAR) were measured following sequential injections of glucose (10 mM), oligomycin (1 μg/mL), and 2-DG (100 mM) at final concentrations.

### Low glucose medium

DMEM without glucose (Thermo Fisher Scientific, 11966025), was used to prepare 10 mM or 0.15 mM glucose DMEM medium, which was then filtered through a 0.2 µm filter. BMDMs were stimulated with or without LPS (100 ng/mL) for 24 hours in 10 mM or 0.15 mM glucose DMEM medium, followed by treatment with ML210 (3 µM) for 12 hours.

### RNA extraction and Real-time PCR

RNA was extracted from lung tissue or cultured cells using RNeasy kit (Qiagen, 74106) according to the manufacturer’s instructions. Complementary DNA was synthesized from 0.5 μg RNA by iScript™ cDNA Synthesis Kit (Bio-Rad, 1708891). Amplification reactions contained a target specific fraction of cDNA and 1 μM forward and reverse primers in iQ™ SYBR® Green Supermix (Bio-Rad, 1708882). Fluorescence was monitored and analyzed in a CFX connect real-time PCR system (Bio-Rad). Gene expression was normalized to β-actin using the delta delta cycle threshold method. Amplification of specific transcripts was confirmed by melting curve analysis at the end of each PCR experiment. The primers used are as follows: Human MCP-1 (Forward: CTGTGCCTGCTGCTCATAGC, Reverse: CAGGTGACTGGGGCATTGATTG); Human IL-1 β(Forward:, Reverse) ; Human VCAM-1(Forward: GGACCACATCTACGCTGACAA, Reverse: AACAGTAAATGGTTTCTCTTGAACA); Human β-actin (Forward: TGTCCCCCAACTTGAGATGT, Reverse: TGTGCACTTTTATTCAACTGGTC); Mouse GCLC (Forward: AGATGATAGAACACGGGAGGAG, Reverse: TGATCCTAAAGCGATTGTTCTTC); Mouse SLC40A1 (Forward: ACCCATCCCCATAGTCTCTGT, Reverse: ACCGTCAAATCAAAGGACCA); Mouse GPX4 (Forward: CCGTCTGAGCCGCTTACTTA a, Reverse: CTGAGAA TTCGTGCATGGAG); Mouse β-actin (Forward: TTCAACACCCCAGCCATGT, Reverse: GTAGATGGGCACAGTGTGGGT);

### Western blot

Proteins were separated by SDS-PAGE through 10% acrylamide gels and transferred to nitrocellulose membranes, blocked with 5% nonfat dry milk in Tween-TBS and reacted with the indicated antibody: GPX4 (Abcam, ab125066) 1:1000 or β-actin (Cell Signaling Technology, 4970) 1:4000 overnight. Membranes were rinsed and incubated with horseradish peroxidase conjugated secondary antibody: Anti-rabbit IgG, HRP-linked Antibody (Cell Signaling Technology, 7074). Reactive proteins were detected by enhanced chemiluminescence, visualized by exposure to radiographic film and quantified by scanning densitometry normalized to β-actin expression measured in each sample on the same gel.

### Statistics

Unless otherwise noted, in vitro experiments were repeated as three independent procedures, with duplicate or triplicate wells averaged prior to statistical analysis. All data were presented as mean ± SD. GraphPad Prism 10.0 was used for statistical analysis. Comparisons between two groups after stimulation were analyzed by two-way ANOVA. LPS dose response experiments in cell cultures were analyzed by one-way ANOVA followed by post hoc T tests using Bonferroni correction for multiple comparisons. P values were indicated as follow: * < 0.05, ** < 0.01, *** < 0.001, **** < 0.0001.

## Acknowledgments

This research received financial support from the New York State Department of Health (C39071GG awarded to B.C.B. and C.G.H.), Pilot grants from the University of Rochester of Department of Medicine (awarded to B.C.B.), and the Department of Environmental Health Sciences (P30 ES001247 awarded to B.C.B. and C.G.H), and Trauma Research and Combat Casualty Care Collaborative (175153 awarded to C.G.H) as well as The University of Texas at San Antonio Startup Funding and The College for Health, Community, and Policy (HCAP) Seed Grant (awarded to C.G.H).

## Author Contributions

C.G.H. designed research; C.G.H., J.Z. performed research; C.G.H., B.C.B. contributed reagents/ analytic tools; C.G.H., J.Z. analyzed data; C.G.H., J.Z., B.C.B. wrote the paper.

